# From DNA-Encoded Library (DEL) Screening to In Vivo Validation: LILRB4 (ILT3)-Targeted Small Molecules Reprograms Myeloid Immune Suppression

**DOI:** 10.64898/2026.06.10.731267

**Authors:** Somaya A. Abdel-Rahman, Moustafa T. Gabr

## Abstract

Alzheimer’s disease (AD) remains a major unmet clinical challenge, with limited therapeutic strategies capable of effectively modulating neuroimmune dysfunction. Leukocyte immunoglobulin-like receptor B4 (LILRB4/ILT3) has recently emerged as an inhibitory microglial immune checkpoint implicated in ApoE-mediated suppression of amyloid-β (Aβ) clearance and inflammatory signaling, supporting its potential as a therapeutic target in AD. Here, we applied DNA-encoded library (DEL) screening of approximately 3.6 billion compounds to identify small molecule binders of LILRB4. Biophysical validation identified **APX1** as a direct LILRB4 ligand with submicromolar affinity, which was further confirmed by cellular thermal shift assay (CETSA). Docking-guided mutagenesis studies defined a discrete ligand-binding interface involving key hotspot residues required for stable target engagement. Functionally, **APX1** disrupted the LILRB4-ApoE interaction in orthogonal ELISA and biolayer interferometry assays. In human iPSC-derived microglia, **APX1** suppressed SHP1/2 phosphorylation, attenuated NF-κB activation and IL-1β secretion, and restored Aβ42 uptake under ApoE-driven inflammatory conditions. **APX1** further demonstrated favorable in vitro developability, metabolic stability, and CNS exposure properties. In the 5xFAD mouse model of AD, oral administration of **APX1** improved cognitive performance, reduced cortical and hippocampal Aβ42 burden, suppressed neuroinflammatory cytokines, and decreased activated microglial populations. Collectively, these findings establish **APX1** as a promising small molecule modulator of the LILRB4-ApoE signaling axis and support pharmacological targeting of neuroimmune checkpoints as a therapeutic strategy for AD.

## INTRODUCTION

Alzheimer’s disease (AD) continues to represent a substantial unmet medical need, as currently approved disease-modifying interventions are primarily restricted to amyloid-directed antibodies that show only moderate clinical efficacy and are further challenged by safety liabilities and inadequate penetration into the central nervous system (1–5). Mounting evidence has positioned microglia as pivotal mediators of amyloid-β (Aβ) pathology, with disease progression increasingly linked to dysregulated signaling networks that govern the balance between activating and inhibitory immune pathways within these cells (2,6–10). Microglial activity is tightly controlled by opposing receptor systems, including activating receptors such as triggering receptor expressed on myeloid cells 2 (TREM2) and inhibitory receptors such as CD33 and members of the leukocyte immunoglobulin-like receptor (LILR) family, which together regulate inflammatory responses and phagocytic function (4,5). Recent reports, including studies from our group, have further demonstrated that pharmacological modulation of TREM2 using small molecules can reshape microglial phenotypes, highlighting the therapeutic potential of targeting neuroimmune checkpoints in AD (11–14).

Among these inhibitory pathways, the immune receptor leukocyte immunoglobulin-like receptor B4 (LILRB4, ILT3) has recently gained attention as a critical microglial checkpoint that negatively regulates Aβ clearance through ApoE-associated signaling mechanisms (15–22). Elevated expression of LILRB4 has been observed in microglia derived from patients with AD, particularly within plaque-associated microglial populations, where receptor levels correlate with ApoE expression and pathological hallmarks such as phosphorylated tau accumulation (23). At the mechanistic level, activation of LILRB4 initiates signaling through immunoreceptor tyrosine-based inhibitory motifs (ITIMs), leading to recruitment of the SHP1 and SHP2 phosphatases and subsequent suppression of cytoskeletal dynamics and phagocytic programs necessary for efficient amyloid removal. Notably, antibody-based inhibition of LILRB4 has been shown to restore microglial activation states, enhance Aβ uptake, and reduce amyloid plaque burden by nearly 50% in preclinical models, while simultaneously dampening interferon-associated transcriptional programs and remodeling plaque-associated microglial populations (23). Collectively, these findings support LILRB4 as an attractive therapeutic target in AD and identify the LILRB4-ApoE axis as a pharmacologically tractable interface for therapeutic intervention.

DNA-encoded library (DEL) screening has become a powerful platform for discovering small molecules capable of reversibly binding protein targets, enabling exploration of chemical diversity far beyond what is achievable using conventional high-throughput screening (HTS) methods (24–32). Continuous improvements in DEL synthesis chemistry, encoding strategies, affinity-selection workflows, library architecture, and sequencing-based analytical pipelines have substantially increased the versatility and scalability of DEL technologies, leading to their widespread integration into modern pharmaceutical and academic drug discovery programs (26–32). Importantly, DEL approaches have shown particular value for identifying ligands against difficult target classes, including protein-protein interactions (PPIs), membrane-associated proteins, and allosteric regulatory sites that have historically proven challenging for traditional screening campaigns (33–37). DEL systems are composed of DNA-barcoded small molecules, where each chemical entity is covalently conjugated to a unique oligonucleotide sequence encoding its synthetic origin (24–26). Library generation generally employs iterative split-and-pool synthetic cycles, in which sequential chemical reactions are accompanied by progressive extension of DNA barcodes (24,25) This highly modular strategy enables the rapid assembly of extremely large combinatorial collections; for instance, incorporation of roughly 1,000 building blocks across three rounds of synthesis can theoretically produce approximately 10^9^ distinct compounds. Such libraries support pooled affinity-selection experiments requiring only limited amounts of purified protein target, thereby significantly enhancing screening throughput and reducing resource demands compared with classical HTS methodologies (24,25). During affinity selection, DEL libraries are exposed to immobilized target proteins, allowing selective enrichment of interacting compounds while nonbinding molecules are eliminated through stringent washing procedures. Bound ligands are then decoded through next-generation DNA sequencing, where the DNA tags serve as molecular barcodes that enable rapid identification of enriched binders from extraordinarily large chemical collections (24,25)

In this work, we utilized DEL screening to discover small molecule ligands targeting LILRB4 (ILT3), followed by comprehensive biophysical characterization, functional evaluation, and in vivo validation. Our findings provide a tractable small molecule framework for pharmacological modulation of the LILRB4/ApoE signaling pathway and address an important unmet challenge in targeting this signaling axis therapeutically.

## RESULTS

### Discovery of LILRB4-targeted small molecules using DEL screening

DEL-based affinity screening against LILRB4 was carried out based on our previously reported procedure (38) using approximately 3.6 billion structurally diverse compounds, allowing broad interrogation of chemical space through pooled selection experiments. The screening campaign involved three sequential rounds of affinity enrichment in which His-tagged LILRB4 protein was immobilized on Ni-NTA magnetic beads and incubated with the pooled DEL mixture, followed by stringent washing steps to eliminate noninteracting molecules and thermal elution of retained binders (Figure 1A). Parallel no-target control (NTC) selections were conducted under identical experimental conditions in the absence of LILRB4 protein to identify compounds enriched through nonspecific interactions. After completion of the third selection cycle, recovered DNA barcodes were amplified by PCR and analyzed by next-generation sequencing (NGS) to determine the identity and abundance of enriched compounds.

**Figure 1.**
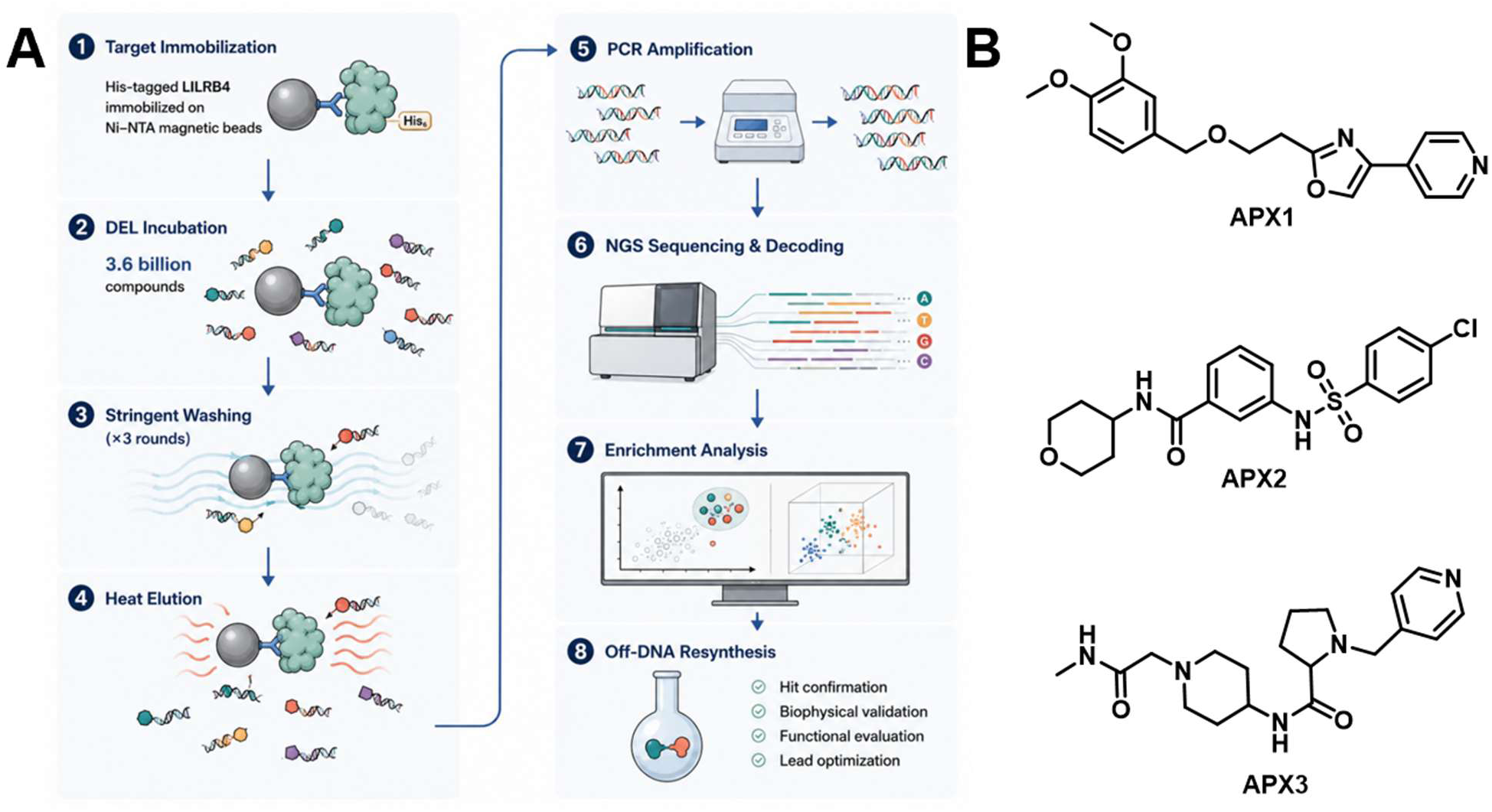
DEL-based discovery of LILRB4-interacting small molecules. **(A)** Illustration of the DEL screening strategy used to identify LILRB4 binders. His-tagged LILRB4 protein was captured on Ni-NTA magnetic beads and exposed to a 3.6 billion-compound DEL. Following incubation, multiple rounds of stringent washing were carried out to eliminate nonspecific binders, and retained molecules were recovered through heat-mediated elution. The associated DNA tags were subsequently amplified by PCR and analyzed by next-generation sequencing (NGS). Sequencing outputs were translated into corresponding chemical structures, and enrichment was determined by comparing target-selection counts against no-target control samples. Candidate hits showing selective enrichment were synthesized off-DNA and advanced for downstream biophysical characterization. **(B)** Chemical structures of three representative hit compounds obtained from the DEL campaign. **APX1**, **APX2**, and **APX3** correspond to structurally distinct chemotypes prioritized for follow-up studies based on their preferential enrichment in the LILRB4-targeted selection assay.

Sequencing outputs were subsequently processed, which converted raw FASTQ sequencing files into corresponding chemical structures with associated copy number information for each experimental condition. Compound enrichment was determined by comparing sequencing counts obtained from LILRB4 selections relative to NTC samples, and the resulting datasets were analyzed and visualized using the DataWarrior platform. In the resulting two-dimensional and three-dimensional enrichment maps, each data point corresponded to an individual DEL member, with axes representing the associated building block combinations. This analysis identified a relatively small subset of compounds showing pronounced enrichment over no-target controls. These candidates demonstrated reproducible enrichment trends across selection rounds together with clear structure-enrichment relationships, supporting specific engagement with LILRB4 rather than nonspecific retention. Based on these criteria, three chemically distinct hit compounds (**APX1**-**APX3**) were selected for off-DNA synthesis and downstream experimental characterization (Figure 1B). The emergence of multiple independent chemotypes from a single DEL campaign underscores the robustness of the screening workflow and supports the existence of tractable small molecule binding sites on LILRB4.

### Orthogonal validation of LILRB4 interaction

The three prioritized hit compounds (**APX1-APX3**) were subsequently evaluated using microscale thermophoresis (MST) with recombinant human LILRB4 extracellular domain protein to validate direct target engagement. All three compounds demonstrated reproducible concentration-dependent binding responses consistent with specific interaction with LILRB4. Among the tested compounds, **APX1** emerged as the highest-affinity binder, exhibiting a dissociation constant (K_D_) of 294 ± 15.9 nM (Figure 2A). In contrast, **APX2** and **APX3** displayed substantially weaker binding affinities, with K_D_ values of 14.5 ± 2.3 μM and 26.4 ± 4.5 μM, respectively. The MST binding curve for **APX1** showed clear saturation behavior at higher ligand concentrations together with stable fluorescence baselines across replicates, supporting specific target engagement rather than nonspecific aggregation-associated effects.

**Figure 2.**
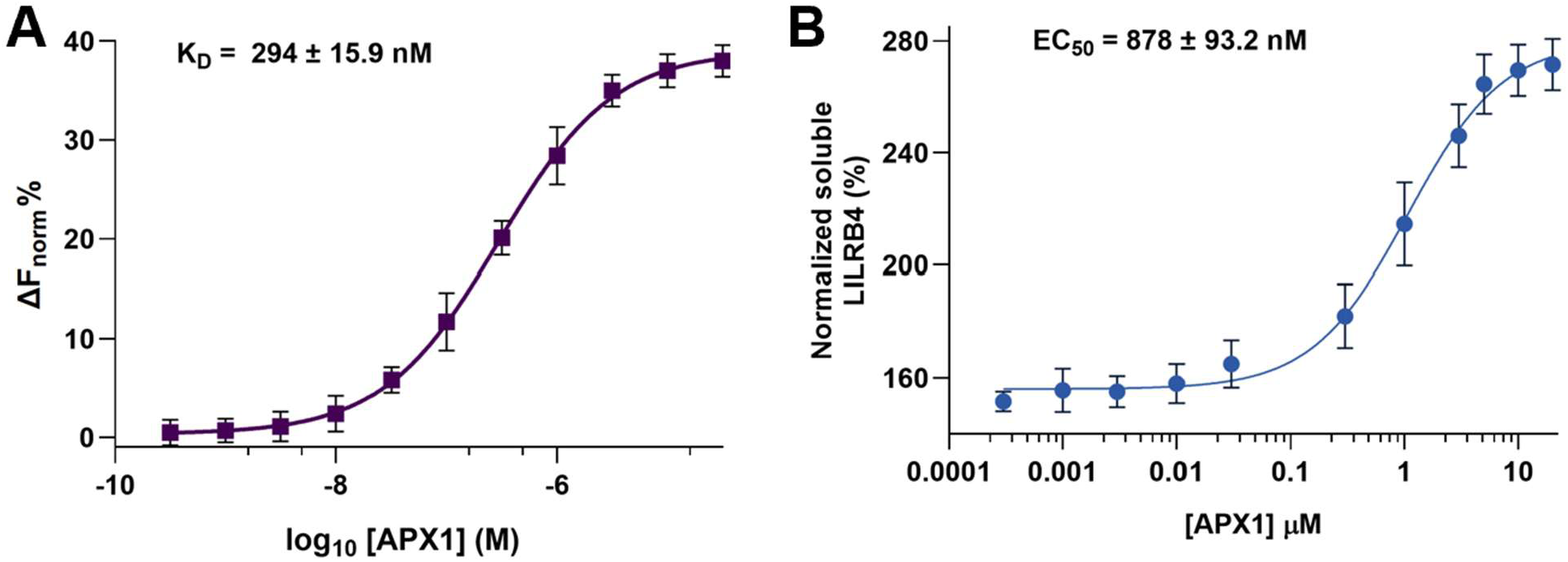
Biophysical and cellular assays confirm direct engagement of LILRB4 by APX1. **(A)** Microscale thermophoresis (MST) analysis demonstrating direct binding of **APX1** to the recombinant extracellular domain of human LILRB4. Values are presented as mean ± SD (n = 5). **(B)** Cellular thermal shift assay (CETSA) showing concentration-dependent stabilization of LILRB4 upon treatment with **APX1**. Cells expressing LILRB4 were incubated with increasing concentrations of the compound prior to thermal challenge, followed by measurement of soluble LILRB4 protein levels. Data are shown as mean ± SD (n = 5).

Given its superior affinity profile, **APX1** was prioritized for downstream validation and mechanistic studies. To further confirm direct target engagement in a cellular context, **APX1** was evaluated using a cellular thermal shift assay (CETSA) adapted from our optimized LILRB4 CETSA platform. Cells expressing LILRB4 were treated with increasing concentrations of **APX1** followed by thermal challenge and quantification of soluble LILRB4 protein. **APX1** induced concentration-dependent thermal stabilization of LILRB4 with an EC_50_ value of 878 ± 93.2 nM, consistent with direct intracellular target engagement and supporting retention of binding activity in a native cellular environment (Figure 2B). Therefore, our results establish **APX1** as a direct small molecule binder of LILRB4 and support its advancement for downstream functional and mechanistic characterization studies.

To gain structural insight into the interaction between **APX1** and LILRB4, molecular docking studies were performed using the extracellular domain of LILRB4. As illustrated in Figure 3A, docking analysis predicted that **APX1** occupies a discrete surface-accessible pocket within LILRB4 and adopts a favorable binding orientation stabilized through a combination of hydrogen bonding, hydrophobic, and aromatic interactions. The predicted docking pose positioned **APX1** in close proximity to residues A70, H180, Y181, and L182, which collectively formed a compact interaction network surrounding the ligand. Among these residues, H180 and Y181 appeared centrally involved in stabilizing APX-01, whereas A70 and L182 contributed additional peripheral interactions within the predicted binding interface.

**Figure 3.**
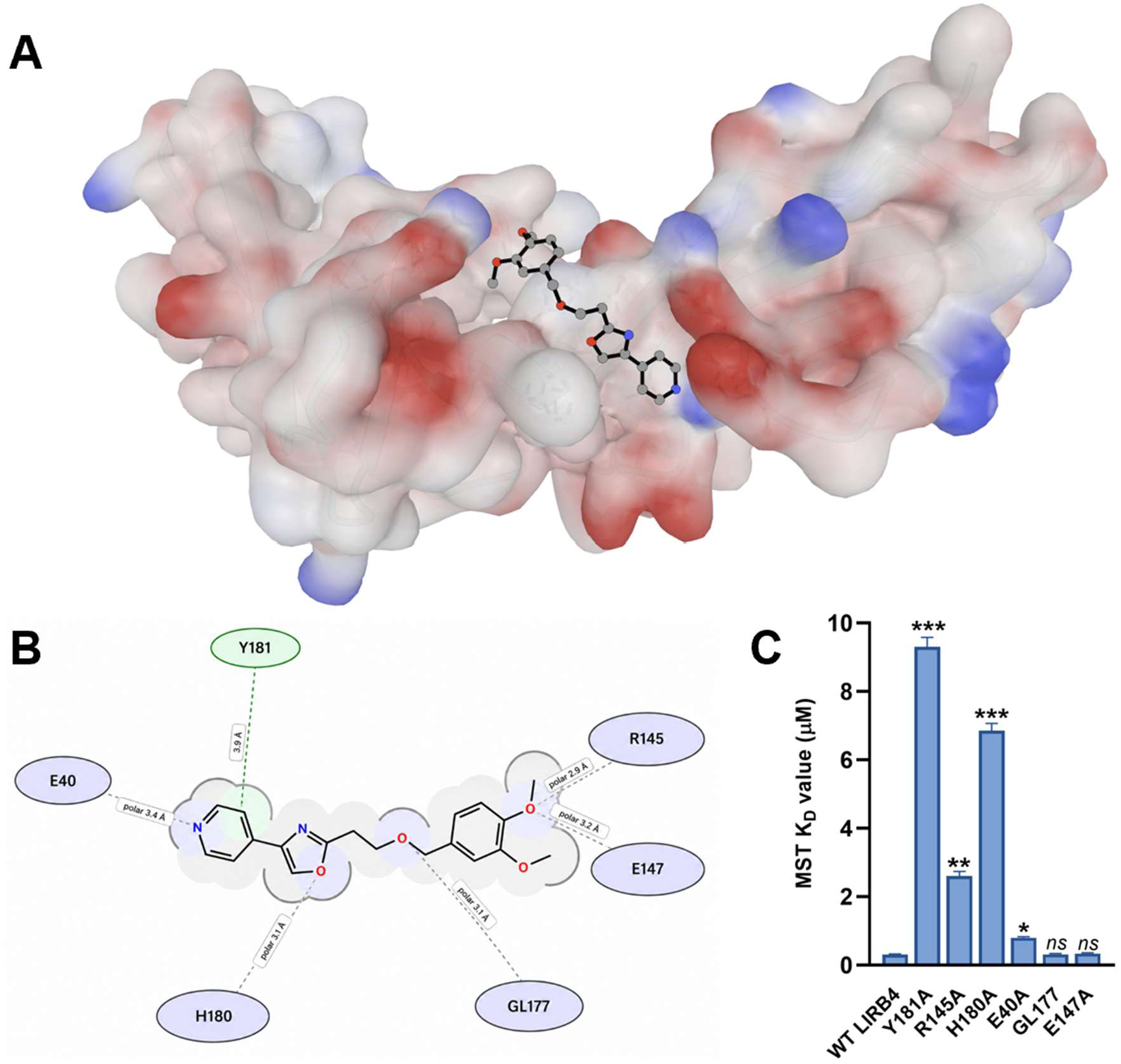
Docking-guided binding mode analysis and site-directed mutagenesis validation of APX1 interaction with LILRB4. **(A)** Predicted three-dimensional binding pose of **APX1** within the LILRB4 binding pocket shown on the surface representation of the receptor. **APX1** occupies a surface-accessible groove enriched in residues implicated in ligand recognition. The protein surface is colored according to hydrophobicity. **(B)** Two-dimensional interaction map of the predicted **APX1**-LILRB4 complex highlighting hydrogen bonding, hydrophobic, and aromatic interactions stabilizing ligand binding. **(C)** Site-directed mutagenesis analysis of residues predicted to contribute to **APX1** binding. Recombinant LILRB4 mutant proteins were evaluated by MST, and K_D_ values were compared relative to wild-type (WT) LILRB4. Mutations of key hotspot residues substantially impaired **APX1** binding, whereas secondary interface mutations produced more moderate effects, supporting the docking-predicted interaction model. Data represent mean ± SD (n = 5). Statistical significance was determined relative to WT LILRB4. \**p* < 0.05, \*\**p* < 0.01, \*\*\**p* < 0.001, and *ns* denotes non-significant.

To further characterize the residue-level interaction landscape, a two-dimensional interaction map was generated from the docked **APX1**-LILRB4 complex (Figure 3B). This analysis revealed multiple predicted interactions supporting the proposed docking orientation. Specifically, hydrogen bonding interactions were observed with A70 and H180, while hydrophobic contacts were identified with A70 and L182. In addition, **APX1** formed π–π stacking interactions with Y181 and H180, including two distinct hydrogen bonding interactions involving H180. Together, these computational analyses suggested that H180 and Y181 function as central hotspot residues mediating **APX1** recognition, whereas A70 and L182 contribute secondary stabilizing contacts that help orient the ligand within the binding pocket.

To experimentally validate the proposed binding model, site-directed mutagenesis followed by MST analysis was performed using alanine-substituted LILRB4 variants (Figure 3C). Wild-type LILRB4 displayed high-affinity interaction with **APX1**, whereas mutations within the predicted interaction interface resulted in varying degrees of affinity reduction. Notably, H180A and Y181A produced the most substantial impairments in **APX1** binding and exhibited highly significant reductions in affinity (\*\*\**p* < 0.001), indicating that these residues serve as critical hotspot interactions required for stable ligand recognition. In contrast, mutation of L182A resulted in a more moderate but still significant reduction in affinity (\*\**p* < 0.01), consistent with a supportive stabilizing role within the binding interface. Mutation of A70A produced a smaller but statistically significant effect on **APX1** binding (\**p* < 0.05), suggesting a more peripheral contribution to ligand stabilization. Additional mutations outside the core interaction interface displayed minimal or nonsignificant effects, further supporting the specificity of the docking-predicted binding pocket. Collectively, the combined docking and mutagenesis analyses establish a structurally coherent model for **APX1** engagement of LILRB4 and provide mechanistic insight into the molecular determinants governing ligand recognition. The pronounced effects associated with H180A and Y181A support the conclusion that these residues function as primary anchoring interactions required for stable complex formation, whereas L182 and A70 contribute secondary stabilizing contacts that help orient **APX1** within the binding pocket. These findings further support the conclusion that **APX1** engages LILRB4 through a discrete and target-specific binding interface and provide a structural framework for future optimization of LILRB4-directed small molecules.

### Functional disruption of the LILRB4-ApoE interaction by APX1

Following confirmation of direct **APX1** engagement with LILRB4 through biophysical and mutagenesis studies, we next investigated whether ligand binding translated into functional inhibition of the biologically relevant LILRB4-ApoE interaction. Because ApoE has been identified as a key endogenous ligand of LILRB4 implicated in suppressive microglial signaling, we evaluated the ability of **APX1** to disrupt this protein-protein interaction using complementary orthogonal assay platforms.

We first employed a plate-based ELISA competition assay in which recombinant LILRB4 extracellular domain protein was immobilized and incubated with ApoE in the presence of increasing concentrations of **APX1**. As shown in Figure 4A, **APX1** produced a robust concentration-dependent reduction in ApoE binding, generating a well-defined sigmoidal inhibition curve consistent with competitive disruption of the interaction interface (IC_50_ value of 375 ± 28.1 nM). At higher concentrations, **APX1** achieved near-complete suppression of ApoE association with LILRB4, supporting strong inhibitory activity against the LILRB4-ApoE complex (Figure 4A).

**Figure 4.**
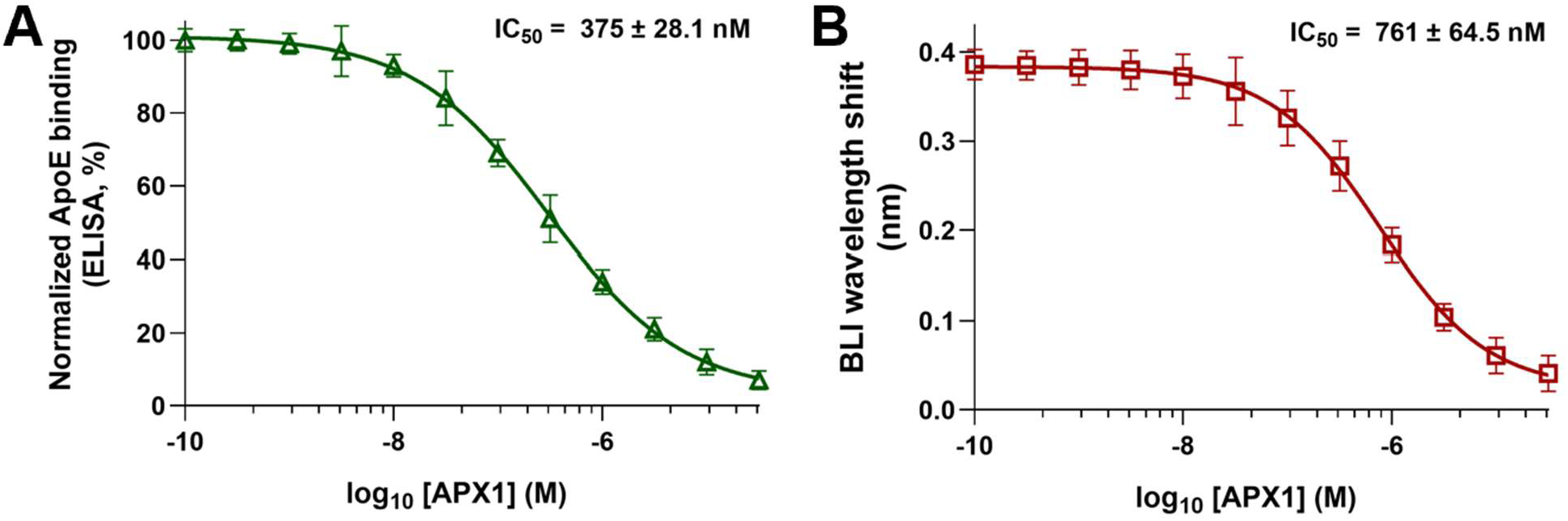
Orthogonal biochemical validation of APX1-mediated disruption of the LILRB4-ApoE interaction. **(A)** Dose-dependent inhibition of ApoE binding to LILRB4 by **APX1** measured by ELISA. Data are presented as normalized ApoE binding (%) and shown as mean ± SD (n = 5). **(B)** BLI-based analysis of **APX1**-mediated inhibition of the LILRB4-ApoE interaction demonstrating a concentration-dependent reduction in binding response. Data are shown as mean ± SD (n = 5).

To further validate these findings using an independent real-time biophysical approach, competitive binding studies were subsequently performed by biolayer interferometry (BLI). In this assay format, immobilized LILRB4 was exposed to ApoE in the presence of increasing concentrations of **APX1**, enabling continuous monitoring of binding responses. Consistent with the ELISA results, **APX1** induced a concentration-dependent decrease in ApoE association with LILRB4 (IC_50_ value of 761 ± 64.5 nM), supporting direct functional inhibition of the interaction interface (Figure 4B). The strong concordance observed between ELISA and BLI across two orthogonal assay systems provides compelling evidence that **APX1** directly disrupts the LILRB4-ApoE interaction rather than indirectly modulating assay output. Together, these results demonstrate that **APX1** not only binds LILRB4 with submicromolar affinity but also effectively converts this interaction into functional blockade of a biologically relevant immune checkpoint pathway. The combined structural, biophysical, and biochemical findings establish a clear mechanistic relationship between target engagement and inhibition of the LILRB4-ApoE signaling axis and provide a strong foundation for downstream cellular and therapeutic evaluation of **APX1**.

### APX1 reverses ApoE-mediated suppressive signaling and restores microglial function in human iPSC-derived microglia

To investigate whether inhibition of the LILRB4-ApoE signaling axis by **APX1** produces functional effects in disease-relevant cellular systems, we evaluated **APX1** in human iPSC-derived microglia exposed to Aβ42-driven inflammatory conditions. Microglial cultures were stimulated with Aβ42 oligomers in the presence or absence of ApoE to mimic ligand-engaged LILRB4 signaling and subsequently treated with **APX1** at increasing concentrations (1, 5, and 10 μM).

We first examined proximal downstream signaling events associated with LILRB4 activation by measuring phosphorylation of the inhibitory phosphatases SHP1 and SHP2, which are recruited to LILRB4 ITIM domains following receptor engagement. Co-treatment with Aβ42 and ApoE induced a pronounced increase in phospho-SHP1 and phospho-SHP2 levels compared with either vehicle-treated cells or Aβ42 stimulation alone, consistent with activation of suppressive LILRB4 signaling pathways. **APX1** treatment produced a concentration-dependent reduction in SHP1 and SHP2 phosphorylation, with partial inhibition observed at 1 μM and more pronounced suppression at 5 and 10 μM, supporting effective blockade of ApoE-induced LILRB4 signaling in human microglia (Figure 5A,B).

**Figure 5.**
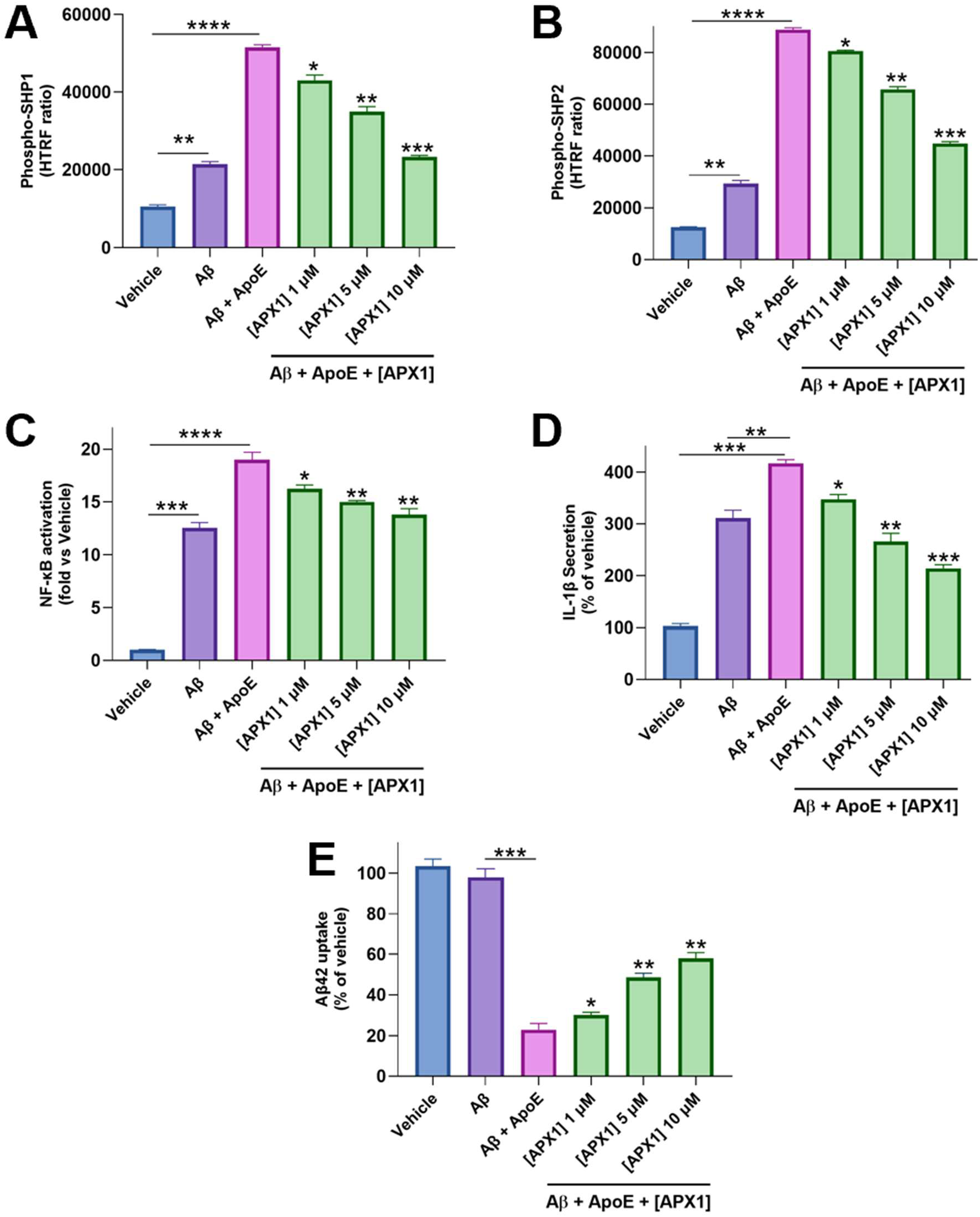
APX1 suppresses LILRB4-dependent signaling, attenuates neuroinflammatory responses, and restores amyloid-β uptake. (A,B) Phosphorylation of SHP1 **(A)** and SHP2 **(B)** following stimulation with Aβ42 and ApoE. Co-treatment with ApoE markedly increased SHP1/2 phosphorylation relative to Aβ42 alone, consistent with activation of LILRB4-dependent inhibitory signaling. **APX1** reduced phospho-SHP1 and phospho-SHP2 levels in a concentration-dependent manner. **(C)** NF-κB activation (fold change relative to vehicle). Aβ42 stimulation increased NF-κB activation, which was further enhanced by ApoE co-treatment. **APX1** suppressed NF-κB signaling in a dose-dependent manner. **(D)** IL-1β secretion (% of vehicle). Aβ42 increased IL-1β release, and this effect was further potentiated by ApoE. **APX1** significantly reduced IL-1β secretion across all tested concentrations. **(E)** Aβ42 uptake (% of vehicle). Aβ42 alone did not significantly alter microglial uptake, whereas co-treatment with ApoE markedly impaired Aβ42 internalization. **APX1** restored Aβ42 uptake in a dose-dependent manner. Statistical significance was determined by one-way ANOVA with appropriate multiple-comparisons tests. Comparisons between Aβ42 and vehicle controls, as well as between Aβ42 + ApoE and corresponding treatment groups, are indicated where appropriate. For **APX1**-treated groups, statistical comparisons were performed relative to the Aβ42 + ApoE condition unless otherwise specified. \*\**p* < 0.01, \*\*\**p* < 0.001, \*\*\*\**p* < 0.0001, and *ns* denotes non-significant.

We next evaluated the impact of **APX1** on downstream inflammatory signaling by quantifying NF-κB activation. Exposure to Aβ42 significantly increased NF-κB activity relative to vehicle-treated controls, and this response was further amplified following ApoE co-treatment (Figure 5C). **APX1** attenuated NF-κB activation in a dose-dependent manner, reducing signaling toward near-basal levels at higher concentrations. The coordinated suppression of SHP1/2 phosphorylation together with reduced NF-κB activation supports a mechanistic connection between LILRB4 engagement and downstream inflammatory transcriptional programs in microglia.

To determine whether modulation of these signaling pathways translated into functional anti-inflammatory effects, secretion of the pro-inflammatory cytokine IL-1β was quantified. Stimulation with Aβ42 elevated IL-1β release, and the addition of ApoE further enhanced cytokine production, consistent with an exacerbated inflammatory phenotype (Figure 5D). **APX1** significantly reduced IL-1β secretion across all tested concentrations, with the strongest suppression observed at 5 and 10 μM, mirroring its inhibitory effects on upstream signaling pathways.

Because impaired amyloid clearance is a central feature of dysfunctional microglial responses in Alzheimer’s disease, we next assessed Aβ42 uptake as a disease-relevant functional readout. Whereas treatment with Aβ42 alone did not substantially alter uptake relative to vehicle controls, co-treatment with ApoE markedly impaired microglial internalization of Aβ42 (Figure 5E), consistent with suppression of phagocytic function through LILRB4 signaling. Importantly, **APX1** restored Aβ42 uptake in a concentration-dependent manner, with significant recovery observed at 5 and 10 μM, indicating functional rescue of microglial phagocytic capacity.

Importantly, **APX1** did not produce significant changes in cell viability under any treatment condition, demonstrating that the observed effects on signaling, cytokine production, and amyloid uptake were not attributable to nonspecific cytotoxicity. Collectively, these findings establish a direct functional relationship between ApoE-mediated LILRB4 activation and suppression of microglial effector functions and further demonstrate that **APX1** effectively converts biochemical target engagement into restoration of disease-relevant microglial activity. Together with the structural, biophysical, and biochemical validation studies, these results support LILRB4 as a tractable neuroimmune checkpoint target and identify **APX1** as a promising small molecule modulator of microglial dysfunction in AD.

### Cross-species target engagement and pharmacokinetic (PK) characterization support in vivo evaluation of APX1 in 5xFAD mice

Before initiating in vivo efficacy studies, we next evaluated the cross-species binding activity and PK properties of **APX1** to determine whether the compound possessed suitable characteristics for animal studies and central nervous system (CNS) exposure. Because subsequent efficacy experiments were performed in murine AD models, we first examined the ability of **APX1** to engage mouse LILRB4 using MST with recombinant mouse LILRB4 extracellular domain protein. **APX1** demonstrated high-affinity binding to murine LILRB4, yielding a K_D_ value of 181 nM (Figure S1), comparable to that observed for the human receptor. These findings indicate that the **APX1** interaction interface is conserved across species and confirm effective cross-species target engagement, thereby supporting the suitability of mouse models for downstream in vivo evaluation.

Preliminary developability and PK characterization of **APX1** revealed a favorable profile across multiple physicochemical, metabolic, and safety-related parameters, supporting its advancement as a small molecule modulator of LILRB4 signaling (Table 1). **APX1** displayed moderate lipophilicity with a LogD_7.4_ value of 3.05, consistent with compounds possessing balanced membrane permeability while avoiding excessive hydrophobicity that can negatively impact solubility or increase nonspecific interactions. Despite containing a heteroaromatic-rich scaffold, **APX1** maintained acceptable aqueous solubility, exhibiting a kinetic solubility of 42.8 μM in 1% DMSO/PBS together with a FaSSIF solubility of 34.5 μM, supporting adequate solubilization under physiologically relevant intestinal conditions (Table 1).

**Table 1.** In vitro PK profile of APX1.

| PK parameter | APX1 |
| --- | --- |
| LogD<sub>7.4</sub><sup>a</sup> | 3.05 |
| Kinetic Solubility<br>1% DMSO/PBS<br>[$\mu\text{M}$]<sup>b</sup> | 42.8 |
| Solubility in<br>FaSSIF [$\mu\text{M}$] | 34.5 |
| PAMPA $P_{app}$<br>[cm/s] | $4.2 \times 10^{-6}$ |
| Stability in<br>simulated gastric<br>fluid (pH 1.2, 2h) | 92.1% |
| Stability in<br>simulated<br>intestinal fluid (pH<br>6.8, 2h) | 88.7% |
| $t_{1/2}$ [min] mouse<br>liver microsomes<sup>c</sup><br>/ CL<sub>int</sub><sup>d</sup> | $68.4 \pm 5.9$ /<br>$26.3 \pm 2.1$ |
| $t_{1/2}$ [min] human<br>liver microsomes<sup>c</sup><br>/ CL<sub>int</sub><sup>d</sup> | $94.6 \pm 7.2$ /<br>$17.9 \pm 1.5$ |
| $t_{1/2}$ [min] mouse plasma | >180 |
| $t_{1/2}$ [min] human plasma | >180 |
| $t_{1/2}$ [min] PBS | >300 |
| PPB human [%]<sup>c</sup> | 94.1 ± 0.8 |
| IC<sub>50</sub> against WI-38 [μM] | >100 |
| IC<sub>50</sub> against HS 27 [μM] | >100 |
| IC<sub>50</sub> against hERG [μM] | >100 |
| CYP3A4 inhibition (% at 10 mM) | 9.2 |
| CYP2D6 inhibition (% at 10 mM) | 5.4 |
| CYP2C9 inhibition (% at 10 mM) | 7.1 |
| CYP2C19 inhibition (% at 10 mM) | 6.0 |

<sup>a</sup>LogD (pH = 7.4) the distribution coefficient at physiological pH 7.4. <sup>b</sup>Kinetic solubility measured in 1% DMSO/PBS (pH 7.4) by UV-vis spectrophotometry (n=3). <sup>c</sup>Values are mean ± SD (n=3). <sup>d</sup>Intrinsic clearance expressed in μL/(min × mg) protein. Values are mean ± SD (n=3).

Evaluation of passive membrane permeability using the PAMPA assay demonstrated a permeability coefficient (P_app_) of 4.2 × 10^−6^ cm/s, suggesting efficient transmembrane diffusion properties. In parallel, **APX1** exhibited strong chemical stability under simulated gastrointestinal conditions, remaining 92.1% intact after 2 h incubation in simulated gastric fluid (pH 1.2) and 88.7% stable in simulated intestinal fluid (pH 6.8). These findings indicate resistance to acid- and intestinal-mediated degradation and support the feasibility of future oral dosing strategies.

Metabolic stability studies further demonstrated that **APX1** possesses favorable microsomal stability in both mouse and human systems. In mouse liver microsomes, **APX1** displayed a half-life of 68.4 ± 5.9 min with an intrinsic clearance (CL_int_) value of 26.3 ± 2.1 μL/(min × mg), while human liver microsome studies revealed improved stability with a half-life of 94.6 ± 7.2 min and a lower intrinsic clearance value of 17.9 ± 1.5 μL/(min × mg), consistent with relatively slow metabolic turnover. **APX1** also demonstrated excellent plasma and buffer stability, remaining stable for more than 180 min in both mouse and human plasma and for over 300 min in PBS, suggesting minimal nonspecific degradation or intrinsic chemical instability (Table 1).

Human plasma protein binding analysis showed high protein association (94.1 ± 0.8%), a profile commonly observed for moderately lipophilic small molecules and one that may contribute to sustained systemic exposure in vivo. Importantly, early safety profiling demonstrated a favorable tolerability profile for **APX1**. No measurable cytotoxicity was observed in WI-38 lung fibroblasts or HS-27 stromal cells at concentrations up to 100 μM. In addition, **APX1** showed no significant inhibition of the hERG potassium channel at concentrations up to 100 μM, suggesting low predicted cardiotoxicity liability. Cytochrome P450 inhibition studies further demonstrated minimal inhibition of major drug-metabolizing isoforms, including CYP3A4 (9.2%), CYP2D6 (5.4%), CYP2C9 (7.1%), and CYP2C19 (6.0%) at 10 μM, supporting a low predicted risk of CYP-mediated drug-drug interactions. These results indicate that **APX1** combines favorable permeability, chemical stability, microsomal stability, and preliminary safety characteristics together with potent LILRB4 target engagement, supporting its continued development and optimization as a neuroimmune-modulating therapeutic candidate.

After intravenous administration into male C57BL/6J mice (8-10 weeks, n=5) at 2 mg/kg, **APX1** exhibited a plasma half-life of 3.4 h, moderate systemic clearance (CL = 10.8 mL/min/kg), and a volume of distribution of 3.2 L/kg, corresponding to a plasma AUC0-∞ of 5.1 µM·h. Following oral dosing at 30 mg/kg, **APX1** achieved a plasma C_max_ of 5.8 µM at T_max_ = 0.75 h, with a plasma AUC0-∞ of 26.4 µM·h, yielding an estimated oral bioavailability of approximately 34%. Importantly, **APX1** also demonstrated robust CNS exposure after oral administration, reaching a brain C_max_ of 3.9 µM at 1 h with a brain AUC0-∞ of 18.7 µM·h. The resulting brain-to-plasma exposure ratio (K_p_) was approximately 0.71, and correction for predicted unbound fractions yielded an estimated K_p,uu_ of ∼0.52, supporting efficient free-drug equilibration across the blood-brain barrier. Collectively, these PK findings established an exposure profile suitable for evaluating **APX1** in murine models of AD. These results established a coherent exposure profile compatible with evaluation in murine models of AD.

Following confirmation of cross-species target engagement together with favorable PK and CNS exposure properties, we next evaluated the therapeutic efficacy of **APX1** in the 5xFAD transgenic mouse model of AD, which recapitulates key pathological features including amyloid deposition, neuroinflammation, and cognitive impairment. Four-month-old male and female 5xFAD mice together with age-matched wild-type (WT) littermates were randomized into treatment groups (n = 12 per group; 6 males and 6 females) and treated with **APX1** (30 mg/kg, PO, QD) or vehicle for 21 consecutive days. This age was selected because 5xFAD mice at this stage display robust amyloid pathology and neuroinflammatory changes while remaining responsive to therapeutic intervention. Behavioral testing was conducted during the final week of treatment, followed by tissue collection for biochemical and immunological analyses.

To assess cognitive function, hippocampal-dependent working memory was evaluated using the Y-maze spontaneous alternation assay (Figure 6A). Vehicle-treated 5xFAD mice displayed significantly impaired alternation performance relative to WT controls, consistent with cognitive deficits characteristic of this model. Treatment with **APX1** significantly improved spontaneous alternation behavior in 5xFAD mice, restoring performance toward WT levels, while producing no detectable effect in WT animals. Importantly, total arm entries did not differ significantly among experimental groups (Figure 6B), indicating that the observed behavioral improvements were not attributable to altered locomotor activity.

**Figure 6.**
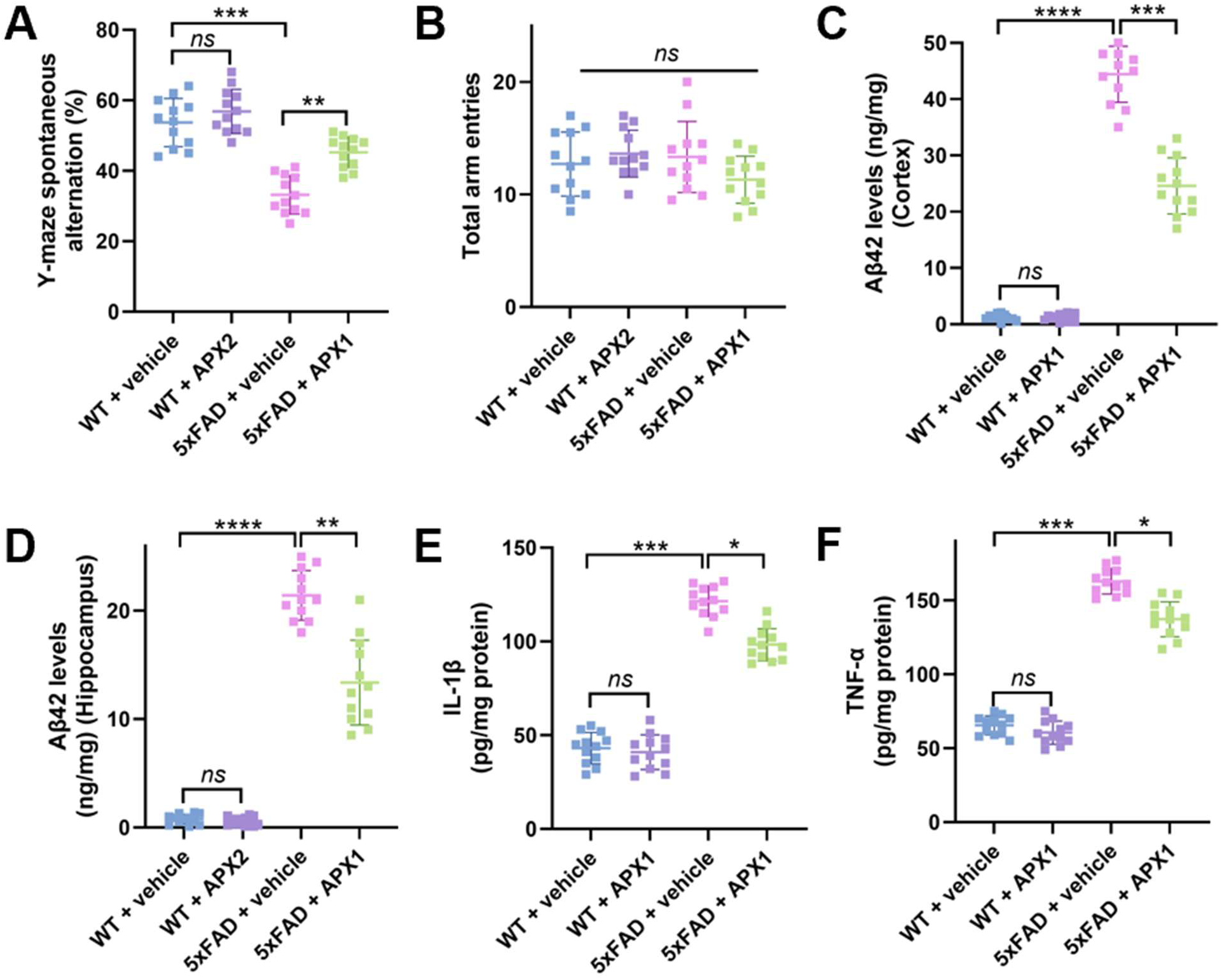
In vivo efficacy of APX1 in the 5xFAD mouse model of AD. **(A)** Y-maze spontaneous alternation performance. Vehicle-treated 5xFAD mice exhibited impaired working memory relative to WT controls, whereas **APX1** treatment significantly improved alternation behavior. **(B)** Total arm entries in the Y-maze showing no significant differences among groups, indicating no effect on locomotor activity. **(C,D)** Aβ42 levels in cortex **(C)** and hippocampus **(D)**. 5xFAD mice displayed elevated amyloid burden compared with WT controls, which was significantly reduced following **APX1** treatment. **(E,F)** Brain levels of pro-inflammatory cytokines IL-1β **(E)** and TNF-α **(F)**. Cytokine levels were elevated in vehicle-treated 5xFAD mice and attenuated following **APX1** administration. Data are presented as mean ± SD (n = 12 per group). Statistical significance was determined using one-way ANOVA with appropriate post hoc multiple-comparisons tests. \**p* < 0.05, \*\**p* < 0.01, \*\*\**p* < 0.001, \*\*\*\**p* < 0.0001, and *ns* denotes non-significant.

We next investigated whether the cognitive improvements observed following **APX1** treatment were associated with modulation of amyloid pathology. Quantification of Aβ42 levels demonstrated marked accumulation in both cortical and hippocampal tissues of vehicle-treated 5xFAD mice relative to WT controls (Figure 6C,D). Treatment with **APX1** significantly reduced Aβ42 burden in both brain regions, producing approximately 40% reduction in cortex and 35% reduction in hippocampus. Because amyloid pathology is closely linked to neuroinflammatory signaling, we additionally evaluated inflammatory cytokine production in brain tissue. Vehicle-treated 5xFAD mice exhibited elevated levels of IL-1β and TNF-α, whereas **APX1** treatment significantly attenuated both inflammatory mediators (Figure 6E,F), consistent with suppression of neuroinflammatory signaling in vivo.

To further assess the impact of **APX1** on microglial activation states, flow cytometric analysis of CD86⁺ microglia was performed as a marker of pro-inflammatory activation. Vehicle-treated 5xFAD mice exhibited a substantial increase in CD86⁺ activated microglia relative to WT controls, whereas **APX1** treatment significantly reduced this population (Figure S2), supporting a shift toward a less inflammatory microglial phenotype. Taken together, these findings demonstrate that **APX1** improves cognitive performance, reduces amyloid burden, suppresses neuroinflammatory signaling, and attenuates pathological microglial activation in the 5xFAD model without affecting general locomotor behavior. Collectively, these results support **APX1** as a promising disease-modifying small-molecule therapeutic targeting neuroimmune dysfunction in AD.

In summary, this study supports LILRB4 (ILT3) as a tractable neuroimmune checkpoint target for small molecule therapeutic intervention in AD. Using DEL screening combined with orthogonal biophysical, structural, cellular, and in vivo validation approaches, we identified **APX1** as a direct LILRB4 binder capable of disrupting the LILRB4-ApoE signaling axis. **APX1** restored microglial phagocytic activity, suppressed inflammatory signaling, demonstrated favorable CNS exposure properties, and reduced amyloid pathology and neuroinflammation in the 5xFAD model. Collectively, these findings support pharmacological modulation of LILRB4 as a promising strategy for reprogramming dysfunctional microglial responses in AD and provide a foundation for the continued development of neuroimmune checkpoint-targeted small molecules.

## EXPERIMENTAL

### DEL screening

DEL screening was performed as we previously reported (38). Prior to DEL screenings, protein immobilization tests were performed to confirm LILRB4 can bind to the magnetic beads through his tag. 10 ul of HispurTM Ni-NTA magnetic beads (Thermo Fisher Scientific, 88832) were incubated with 5 µg of LILRB4 (from SinoBiological, Catalog# 16742-H08H). The composition of the incubation buffer was 1x PBS, 10 mM imidazole, 0.1 mg/ml sssDNA, 0.005% Tween-20, pH 7.4. After 30 min incubation at RT, the supernatant was collected, and the beads were resuspended and washed by 100 µl of the incubation buffer. The supernatant was collected as wash, and the beads were resuspended by 100 µl of the incubation buffer. The bead-suspensions were then heated at 95 °C for 10 min, and the supernatant was collected as heated elution. The heated beads were again resuspended in 100 µl of the incubation buffer. 10 µl of each sample was loaded to a 5%/12% Tris-HCl SDS-PAGE. The gel was run for 150 V for 1 h followed by staining with Coomassie Blue for 1 h and destaining in distilled water overnight.

Approximately 3.6 billion DEL compounds against LILRB4 was performed by panning 10 µg target on 20 µl HispurTM Ni-NTA with the DNA-Encoded Chemical Libraries. Three rounds of affinity screening were performed in each condition, including protein immobilization, DEL panning, wash steps, and heated elution. Heated elution samples from the third round of affinity screening were subjected to the Next-Generation Sequencing to reveal the identity of the bound compounds under each screening condition. The fastq files were converted into chemical identities and the corresponding copies in each screening condition. The data were visualized by DataWarrior program. Each dot in the 2D and 3D plots represents a full molecule with each axis representing the BB index. The molecules with clear structure-enrichment relationship and high copies were considered as promising binders of the target and subjected to off-DNA hit resynthesis and validation.

### Microscale thermophoresis (MST)

Binding affinities of selected hits were quantified by microscale thermophoresis using a Monolith X instrument (NanoTemper Technologies). His-tagged LILRB4 protein (from SinoBiological, Catalog# 16742-H08H) was labeled with RED-tris-NTA 2nd Generation dye using the Monolith His-Tag Labeling Kit (Cat. #MO-L018) following the manufacturer’s guidelines. Compound titrations were prepared as serial dilutions (PBS buffer, pH 7.4, 0.05% Tween-20, 1% (v/v) DMSO).

Following a 15 min incubation at room temperature in the dark, samples were loaded into Monolith capillaries (Cat. #MO-K022) and analyzed at 25 °C using 40% LED power and medium MST power settings. Normalized fluorescence (F_norm_) values were determined as the ratio of fluorescence intensity after and before IR laser heating. Each compound was evaluated in five technical replicates. Dissociation constants (K_D_) were calculated from three independent experiments using MO.Affinity Analysis software and GraphPad Prism 10, applying standard dose-response fitting models. Data represent mean ± SD (n=5).

### ELISA-based competition assay

LILRB4 extracellular domain (2 µg/mL) was immobilized on 96-well high-binding plates (Corning) overnight at 4°C. Plates were blocked with 5% BSA in PBS for 1 h at room temperature. Recombinant human ApoE (R&D Systems) was added at a fixed concentration (100 nM) in the presence of serial dilutions of compounds. Bound ApoE was detected using an HRP-conjugated anti-ApoE antibody, followed by TMB substrate development. Absorbance was measured at 450 nm using a Tecan plate reader. Data were normalized to DMSO controls and fitted to determine IC_50_ values using nonlinear regression (GraphPad Prism v10). Reported values represent mean ± SD (n=5).

### Biolayer interferometry (BLI)

BLI experiments were performed using an Octet RED96 system (Sartorius). LILRB4 protein was immobilized onto Ni-NTA biosensors, followed by equilibration in assay buffer (PBS + 0.05% Tween-20). ApoE (100 nM) was associated in the presence of increasing concentrations of **APX1**. Binding responses were monitored as wavelength shifts, and inhibition curves were generated to determine IC_50_ values. Data were analyzed using Octet Data Analysis software. Reported values represent mean ± SD (n=5).

### CETSA

CHO cells stably expressing human ILT3/LILRB4 (from Creative Biogene, Cat# CSC-RO0667) were maintained in Ham’s F-12 medium supplemented with 10% fetal bovine serum and 1% penicillin-streptomycin under standard humidified culture conditions (37 °C, 5% CO2). Cells were seeded in 384-well plates at 12,000 cells/well and incubated overnight at 37 °C in 5% CO₂. **APX1** was added at the indicated concentrations and incubated for 60 min. Cells were subsequently subjected to thermal challenge at 51 °C for 3 min (using a calibrated Bio-Rad C1000 Touch thermal cycler), followed by cooling to room temperature. Following thermal challenge, cells were lysed directly in assay plates by addition of lysis buffer supplemented with protease inhibitor cocktail and incubated for 30 min at 4 °C with gentle agitation. Lysates were subsequently clarified by centrifugation at 4000 × g for 10 min to remove aggregated and insoluble protein species generated during thermal denaturation.

The soluble fraction of LILRB4 remaining after thermal challenge was quantified using an AlphaLISA-based immunodetection format. Briefly, clarified lysates were incubated with ILT3-specific biotinylated detection antibody and AlphaLISA acceptor beads according to the manufacturer’s instructions, followed by addition of streptavidin-coated donor beads under reduced-light conditions. After incubation at room temperature, AlphaLISA signal was measured using Tecan Spark plate reader. CETSA stabilization signals were normalized to vehicle-treated controls, and dose-response curves were generated using nonlinear regression analysis. Data represent mean ± SD, n = 5.

### Computational studies

Molecular docking studies were performed to characterize the interactions of **APX1** with LILRB4. The crystal structure of LILRB4 (PDB ID: 3P2T) was obtained from the Protein Data Bank. The solvents and water were removed from the receptor prior to docking. The lead compounds in SMILES format were converted into .mol2 and subsequently to .pdbqt using openbabel and MGLTools. The pdbqt file includes Gasteiger charges and information on rotatable bonds. We have carried out molecular docking using AutoDock4.0. The 2D binding pose was generated using Rowansci.com

### Site-directed mutagenesis

Alanine substitutions were introduced into the human LILRB4 extracellular domain using the QuikChange Lightning Site-Directed Mutagenesis Kit (Agilent Technologies) according to the manufacturer’s protocol. Briefly, mutagenic primers were designed to incorporate the desired codon substitutions and PCR amplification was performed using a high-fidelity polymerase. Parental methylated DNA was digested with DpnI, and the resulting plasmids were transformed into *E. coli* DH5α cells. All mutations were confirmed by Sanger sequencing. Recombinant wild-type and mutant LILRB4 proteins were expressed and purified as described above. Protein integrity and purity (>90%) were verified by SDS-PAGE and analytical size-exclusion chromatography. Binding affinities of mutant proteins were evaluated by MST under identical conditions to wild-type controls, and K_D_ values were obtained from nonlinear regression fits as described above.

### Human iPSC-derived microglia assays

Human iPSC-derived microglia (FujiFilm Cellular Dynamics) were cultured in microglia maintenance medium (Axol Bioscience Ltd, Catalog #ax0660) supplemented according to the supplier’s recommendations at 37 °C in a humidified 5% CO_2_ incubator. Cells were plated at 50,000 cells per well in poly-D-lysine-coated 96-well plates and allowed to recover for 24 h prior to treatment.

Aβ_42_ oligomers were prepared by dissolving lyophilized Aβ_42_ peptide (≥95% purity; AnaSpec, Catalog# AS-20276) in HFIP, followed by evaporation, resuspension in DMSO, and oligomerization in PBS at 4 °C for 24 h. Cells were treated with Aβ_42_ (1 μM) in the presence or absence of recombinant human ApoE (100 nM), followed by addition of **APX1** at indicated concentrations (1, 5, and 10 μM). Treatments were performed for 24 h. Vehicle controls contained equivalent DMSO concentrations (<0.5%). Reported values represent mean ± SD (n=5).

### Signaling assays

Phosphorylation of SHP1 and SHP2 was quantified using phospho-specific ELISA kits (abcam, catalog# ab279924 and ab314344, respectively) following the manufacturer’s protocol. Briefly, cells were lysed in ice-cold lysis buffer supplemented with protease and phosphatase inhibitors. Lysates were normalized for total protein content (BCA assay), and equal amounts were loaded per well. Reported values represent mean ± SD (n=5).

NF-κB activation was measured using a luciferase reporter assay in human iPSC-derived microglia. Cells were plated at 50,000 cells per well in white 96-well plates and transfected with an NF-κB firefly luciferase reporter (Promega) and a Renilla luciferase control plasmid (Promega) using Lipofectamine 3000 (Thermo Fisher Scientific, Cat. #L3000008). Twenty-four hours post-transfection, cells were treated with Aβ42 oligomers (1 μM) ± ApoE (100 nM) and **APX1** at indicated concentrations (1, 5, and 10 μM) for 24 h. Luciferase activity was measured using the Dual-Glo Luciferase Assay System (Promega, Cat. #E2920). NF-κB activity was calculated as the ratio of firefly to Renilla signal and expressed as fold change relative to vehicle controls. Reported values represent mean ± SD (n=5).

### Cytokine measurements

IL-1β levels in culture supernatants were quantified using a human IL-1β ELISA kit (R&D Systems, catalog# DLB50). Supernatants were collected, clarified by centrifugation (1,000 × g, 5 min), and analyzed according to the manufacturer’s instructions. Cytokine concentrations were interpolated from standard curves and normalized where appropriate. Reported values represent mean ± SD (n=5).

### Aβ uptake assay

Fluorescently labeled Aβ_42_ (HiLyte Fluor 555-conjugated from AnaSpec, catalog# AS-60480-01) was added to cells at a final concentration of 500 nM following treatment conditions described above. After incubation (4 h), cells were washed extensively with PBS and treated with trypan blue to quench extracellular fluorescence. Intracellular fluorescence was quantified using a Tecan Spark plate reader and normalized to cell number (total protein).

### Cell viability

Cell viability was assessed using the CellTiter-Glo luminescent assay (Promega, catalog#G7570) according to the manufacturer’s protocol. Following treatment, reagent was added directly to wells, incubated for 10 min at room temperature, and luminescence was measured. Viability was normalized to vehicle-treated controls. Reported values represent mean ± SD (n=5).

### PK studies

PK studies were conducted in male C57BL/6 mice (8-10 weeks old; n = 5 per time point). **APX1** was administered by intravenous injection (2 mg/kg) or oral gavage (30 mg/kg) in a formulation consisting of 10% DMSO, 40% PEG400, and 50% sterile saline (v/v/v), prepared fresh prior to dosing. Blood samples were collected at 0.25, 0.5, 1, 2, 4, 8, and 24 h post-dose via tail vein sampling into EDTA-coated tubes. Plasma was isolated by centrifugation at 3,000 × g for 10 min at 4 °C.

For CNS exposure analysis, mice were perfused with ice-cold PBS before brain collection to minimize residual blood contamination. Brain tissues were harvested, weighed, and homogenized in ice-cold PBS. Compound concentrations in plasma and brain homogenates were quantified by LC-MS/MS following protein precipitation with acetonitrile containing an internal standard. Chromatographic separation was performed on a C18 column, and analytes were detected in positive electrospray ionization mode using multiple reaction monitoring. Calibration curves were prepared in blank mouse plasma or brain homogenate matrix and used to calculate compound concentrations. PK parameters were calculated by noncompartmental analysis using Phoenix WinNonlin. Brain-to-plasma exposure ratios were calculated from AUC values. Unbound brain-to-plasma ratios (K_p,uu_) were calculated using measured plasma and brain unbound fractions.

### In vivo efficacy in 5xFAD mice

All procedures were approved by the Institutional Animal Care and Use Committee. Male and female 5xFAD mice and wild-type littermates (4 months old) were randomized into treatment groups (n = 12 per group; 6 males and 6 females per group).

**APX1** was administered by oral gavage at 30 mg/kg once daily (QD) for 21 days. Vehicle-treated animals received formulation only. Treatment allocation, behavioral testing, biochemical assays, flow cytometry analysis, and data quantification were performed by investigators blinded to genotype/treatment.

### Behavioral testing

Cognitive function was assessed using the Y-maze spontaneous alternation assay. Mice were placed in the maze and allowed to explore freely for 8 min. Alternation percentage was calculated as the ratio of sequential entries into all three arms over total possible alternations. Total arm entries were recorded to control for locomotor activity.

### Amyloid quantification

Cortex and hippocampus were microdissected separately, snap-frozen, and homogenized in ice-cold extraction buffer containing protease inhibitors. Tissue homogenates were centrifuged to obtain soluble fractions, and the remaining pellets were further extracted to recover insoluble Aβ. Soluble Aβ was extracted in TBS or PBS-based buffer, and insoluble Aβ was extracted from pellets using 5 M guanidine-HCl or 70% formic acid. Aβ42 levels were quantified using a commercially available ELISA kit (Thermo Fisher Scientific; Aβ42, Cat. # KMB3441), according to the manufacturer’s instructions. Concentrations were interpolated from standard curves and normalized to total protein content determined by BCA assay. Data are reported separately for cortex and hippocampus.

### Cytokine analysis

Brain homogenates were prepared in lysis buffer and cytokine levels (IL-1β, TNF-α) were measured using ELISA kits (R&D Systems, catalog# MLB00C and MTA00B, respectively). Values were normalized to total protein content.

### Microglial activation

Brain tissues (cortex and hippocampus) were rapidly dissected and processed into single-cell suspensions using the Adult Brain Dissociation Kit (Miltenyi Biotec, Cat. #130-107-677) according to the manufacturer’s protocol. Myelin was removed using Myelin Removal Beads II (Miltenyi Biotec, Cat. #130-096-733). Cells were resuspended in FACS buffer (PBS containing 2% fetal bovine serum and 2 mM EDTA) and incubated with anti-mouse CD16/CD32 Fc block (BioLegend, Cat. #101320) for 10 min at 4 °C to prevent non-specific binding.

Cells were stained for 30 min at 4 °C with fluorophore-conjugated antibodies against CD45 (APC, clone 30-F11, BioLegend, Cat. #103112), CD11b (FITC, clone M1/70, BioLegend, Cat. #101206), TMEM119 (PE, clone 106-6, BioLegend, Cat. #157306), and CD86 (PerCP-Cy5.5, clone GL-1, BioLegend, Cat. #105028). Dead cells were excluded using a viability dye (Zombie NIR, BioLegend, Cat. #423106). Data were acquired on a BD LSRFortessa flow cytometer (BD Biosciences) and analyzed using FlowJo v10. Microglia were defined as live CD45^low^ CD11b^+^ TMEM119^+^ cells. Activated microglia were quantified as the percentage of CD86^+^ cells within the microglial population. A minimum of 50,000 live events per sample were recorded.

### Statistical analysis

All data are presented as mean ± SD. Statistical analyses were performed using GraphPad Prism v10. Two-group comparisons were conducted using unpaired two-tailed Student’s *t*-tests. Multiple comparisons were analyzed using one-way ANOVA followed by Tukey’s post hoc test. *P* values < 0.05 were considered statistically significant. Exact sample sizes and statistical tests are indicated in figure legends.

## Data and materials availability

All data needed to evaluate the conclusions in the paper are present in the paper and/or the Supplementary Materials.

