## Supporting Information for "From DNA-Encoded Library (DEL) Screening to In Vivo Validation: LILRB4 (ILT3)-Targeted Small Molecules Reprograms Myeloid Immune Suppression"

| **Contents** | |  |
| --- | --- | --- |
| **APX1** directly binds recombinant murine LILRB4 as determined by MST | | S2 |
| CD86⁺ activated microglia in 5xFAD mice following **APX1** treatment | | S3 |

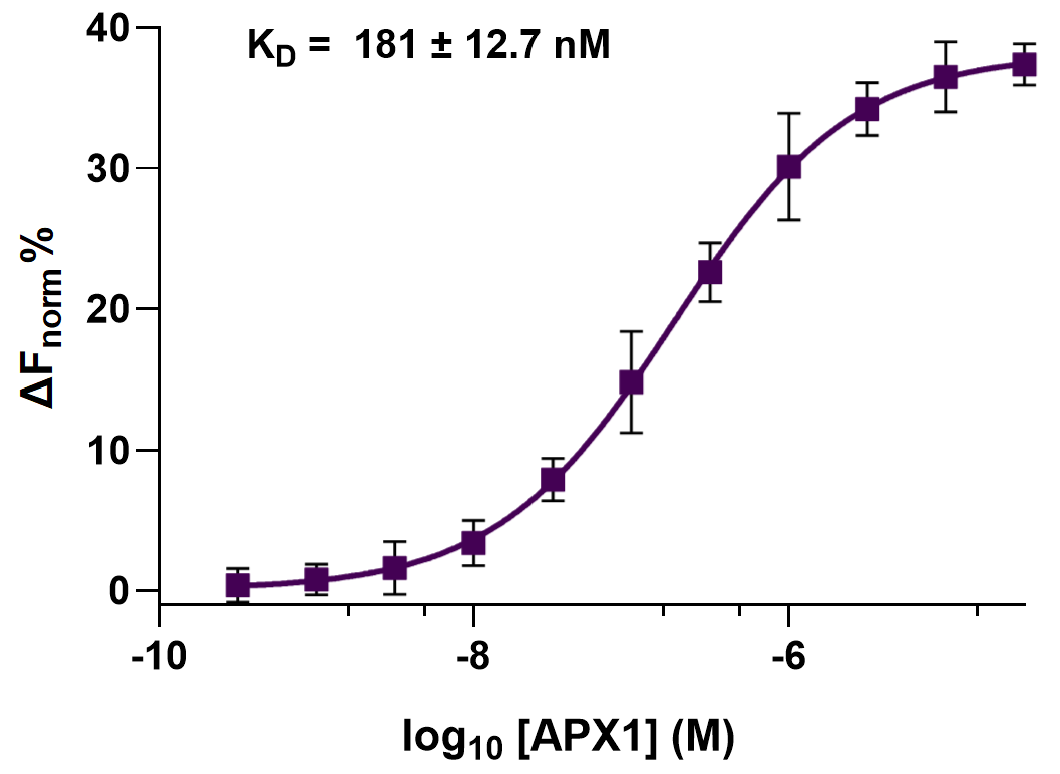

**Figure S1. APX1 directly binds recombinant murine ILT3 as determined by MST.** Binding of **APX1** to recombinant murine LILRB4 was evaluated using MST under solution-phase conditions. **APX1** exhibited concentration-dependent interaction with murine LILRB4. Data represent mean ± SD (n=5).

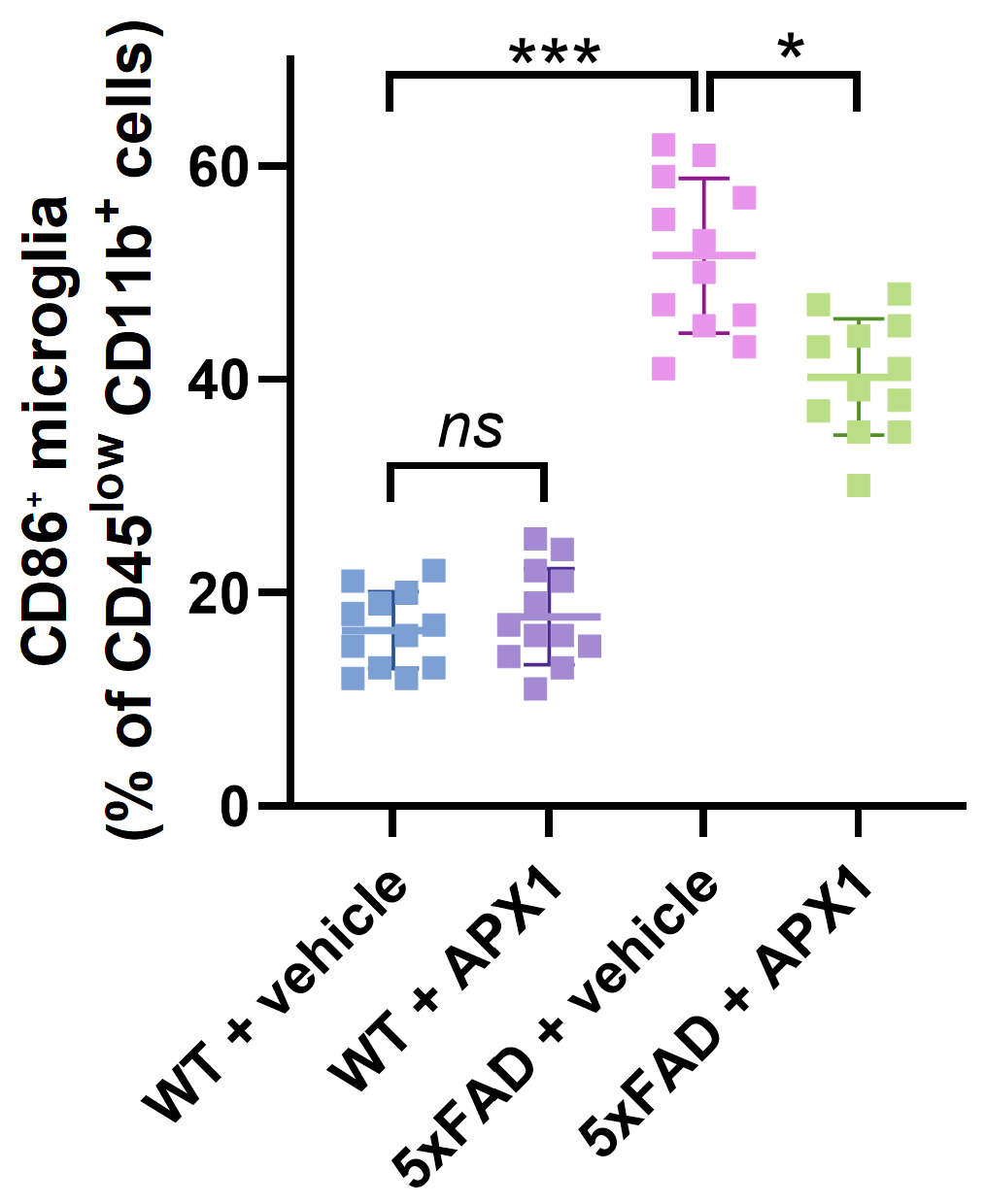

**Figure S2.** CD86⁺ activated microglia (% of CD45^low^ CD11b⁺ microglia). 5xFAD mice showed increased microglial activation that was significantly reduced following **APX1** treatment (30 mg.kg). Data are presented as mean ± SD (n = 12 per group). Statistical significance was determined using one-way ANOVA with appropriate post hoc multiple-comparisons tests. **p* < 0.05, ****p* < 0.001, and *ns* denotes non-significant.
